# A survey of lineage-specific genes in *Triticeae* reveals *de novo* gene evolution from genomic raw material

**DOI:** 10.1101/2022.05.12.491495

**Authors:** Manuel Poretti, Coraline R. Praz, Alexandros G. Sotiropoulos, Thomas Wicker

## Abstract

Plant genomes typically contain ∼35,000 genes, almost all belonging to highly-conserved gene families. Only a small fraction are lineage-specific, which are found in only one or few closely related species. Little is known about how genes arise *de novo* in plant genomes and how often this occurs, however they are believed to be important for plants diversification and adaptation. We developed a pipeline to identify lineage-specific genes in *Triticeae*, using newly available genome assemblies of wheat, barley and rye. Applying a set of stringent criteria, we identified 5,942 candidate *Triticeae*-specific genes (TSGs), of which 2,337 were validated as protein-coding genes in wheat. Differential gene expression analyses revealed that stress-induced wheat TSGs are strongly enriched in secreted proteins. Some were previously described to be involved in *Triticeae* non-host resistance and cold adaptation. Additionally, we show that 1,079 TSGs have sequence homology to transposable elements (TEs), ∼68% of them deriving from regulatory non-coding regions of *Gypsy* retrotransposons. Most importantly, we demonstrate that these TSGs are enriched in transmembrane domains and are among the most highly expressed wheat genes overall. To summarize, we conclude that *de novo* gene formation is relatively rare and that *Triticeae* probably possess ∼779 lineage-specific genes per haploid genome. TSGs which respond to pathogen and environmental stresses, may be interesting candidates for future targeted resistance breeding in *Triticeae*. Finally, we propose that non-coding regions of TEs might provide important genetic raw material for the functional innovation of TM domains and the evolution of novel secreted proteins.

## Introduction

Lineage-specific genes are found only in a single species or in a group of closely related species, and are sometimes also referred to as orphan genes. They are evolutionary novelties that have no sequence homology with any known coding sequences of distantly related species (Tautz & Domazet-Lošo, 2011). Lineage-specific genes can evolve from already existing protein-coding sequences (duplication and divergence, horizontal gene transfer) or arise *de novo* from ancestral non-coding regions. Such *de novo* genes are characterized by short open reading frames (ORFs), low-expression level and are specific to different tissue, developmental stages and stress conditions (Schlötterer, 2015). For these reasons, they are difficult to identify and to distinguish from pseudo-genes or annotation artifacts.

There are two major hypotheses for the evolution of *de novo* genes (Van Oss & Carvunis, 2019). In the “expression-first model”, transcribed non-coding regions (i.e., proto-genes) evolve neutrally and expose genetic variations to selection pressure. Proto-genes might therefore gain adaptive mutations and gradually evolve into more complex and better functional *de novo* genes. On the other hand, the “ORF-first” model postulates that potential ORFs are already present in the genome and become translated after the acquisition of transcriptional factor binding sites. In addition, the “Transmembrane (TM) first” model builds on the expression-first model and proposes that the translation of thymine-rich non-genic regions might favour the emergence of novel adaptive genes that encode proteins with transmembrane domains (Vakirlis *et al*., 2020). Indeed, the authors demonstrated that novel ORFs with beneficial impact on the fitness of yeast, were enriched in TM domains.

Lineage-specific genes represent 10-20% of all annotated genes in sequenced genomes. Even though their function remains mostly unknown, several studies in plants indicated that they may play a role in the adaptation to environmental stresses and in defense responses against pathogens (Khalturin *et al*., 2009; Arendsee *et al*., 2014). An important class of lineage-specific genes involved in plant-pathogen interactions are small, Cysteine-rich and/or Proline-rich and contain a signal peptide or transmembrane domain (Douchkov et al., 2011).

Transposable elements (TEs) make a considerable fraction of all known genomes. They are mobile genetic units that can make copies of themselves and move throughout the genome. TEs are thought to play an important role in the evolution of *de novo* genes, either by providing new promoter elements for already existing ORFs, or by creating new coding-sequences through retroposition, gene capture or exon fusion (Van Oss & Carvunis, 2019). TEs were indeed found to be associated with about half of the identified lineage-specific genes in primates and rice (Toll-Riera *et al*., 2009; Jin *et al*., 2019), suggesting that TE-driven gene evolution might be important in all eukaryotes. However, it is still unclear how frequently *de novo* genes actually emerge.

The study of *de novo* gene formation in large and TE-rich plant genomes is still in its infancy since only recently, chromosome-scale assemblies of the large *Triticeae* genomes of barley, wheat and rye became available (Mascher *et al*., 2017; IWGSC *et al*., 2018; Rabanus-Wallace *et al*., 2021). Such near-complete genome assemblies are necessary if one wants to study how *de novo* genes evolve from repetitive sequences. The release of *Triticeae* genome sequences provides an attractive system to study the evolution of *de novo* genes because it represents a set of species which are closely enough related so that direct comparison of gene content and collinearity is possible, but yet distant enough to have numerous differences.

A number of criteria were defined to assess gene annotation quality of *Triticeae* genomes (Mayer *et al*., 2014; IWGSC *et al*., 2018). High-confidence (HC) genes were defined by significant blast hits across most of their length to proteins found in other reference genomes such as *Brachypodium* or *Arabidopsis*. Low-confidence (LC) genes do not match these criteria and may include fragments of genes and TEs. In *Triticeae*, approximately 34,000-43,000 high-confidence genes were annotated per haploid genome, and similar numbers of low-confidence genes. It is safe to assume that a large fraction of low-confidence genes are annotation artifacts, but they may nevertheless contain some genuine novel genes, especially if they are supported by transcriptome data. However, it is extremely difficult to prove whether a predicted gene without homologs in other species is indeed real because there are usually very few hints as to its function. In protein coding genes, the ratio of substitution rates at non-synonymous and synonymous sites (dN/dS) can be used as an indication that a candidate gene has acquired a function and is therefore under purifying selection. Furthermore, there has to be evidence that the predicted transcript is actually translated into a protein, for example by the presence of matching peptide sequences in proteomics databases.

In this study, we aimed at developing bioinformatics methods to identify candidates for lineage-specific genes in *Triticeae*. In particular, we focused on the identification of genes that originated *de novo*, and we wanted to study how these genes emerged. In total, we identified 5,942 candidate TSGs. In wheat, 2,337 were translated and/or supported by both transcriptomic and negative selection evidence, thus leading us to the conclusion that ∼779 genes per haploid genome evolved *de novo* in *Triticeae*. Approximately 18% (1,079 TSGs) were derived from TE sequences and, in some cases, we could trace their evolution back to specific events such as gene capture. We highlight TE regulatory regions as important hot-spots for the evolution of *de novo* genes, especially for TSGs encoding transmembrane proteins. Finally, we identified a set of genes that encode putative secreted and/or transmembrane proteins that could play a role in the adaptation of *Triticeae* to environmental and pathogenic stresses. We propose that the approach presented here can be used to identify genuine lineage-specific genes, including some interesting candidates that might be selected for future targeted resistance breeding in *Triticeae*.

## Materials and Methods

### Prediction of *de novo* TSGs

Identification of *de novo Triticeae*-specific genes (TSGs) was performed by comparing the previously published high-quality genome annotations (both high-confidence HC and low-confidence LC coding sequences) of three *Triticeae* species, namely wheat (*T. aestivum*, IWGSC RefSeq v1.0; IWGSC *et al*., 2018), rye (*S. cereale*, Lo7 v1; Rabanus-Wallace *et al*., 2021), and barley (*H. Vulgare*, IBSC v2; Mascher *et al*., 2017). Our assumption was that coding sequences that are conserved within wheat, rye, and barley, and that can not be find in other plant species, including closely related ones, might represent potential *de novo Triticeae* genes.

Barley was the first chromosome-scale *Triticeae* genome to be assembled. It was annotated using the PGSB pipeline (initially developed for barley; Mascher *et al*., 2017), whereas the most recent genomes of wheat and rye were annotated using both the PGSB and the TriAnnot (developed for *Triticeae* and initially used for wheat; Leroy *et al*., 2012) pipelines. Most importantly, similar criteria (originally described in Mayer *et al*., 2014 and adapted in IWGSC *et al*., 2018) were used for classifying the annotated coding sequences into HC and LC genes, therefore making our inter-species comparison less biased towards prediction artifacts. Two closely related outgroup species (*Brachypodium distachyon* v3.2 and rice IRGSP v1.0), and the NCBI and UniprotKb databases were considered for filtering out genes which are not specific to the *Triticeae* clade.

Given that automated pipelines generate a large number of erroneous gene predictions, we first checked the integrity of the annotated genes. Only the sequences with an intact open reading frame (ORF; presence of start/stop codons and absence of in-frame stop codons) were considered for further analyses. Then, in order to identify genes that might encode *Triticeae*-specific proteins, we performed Blastp analyses between the protein sequences of *Triticeae* species and the ones of the outgroup species *Brachypodium* and rice. *Triticeae* proteins that had a significant blast hit (evalue 1e-5) with at least one of the two outgroup species were filtered out. Because we are interested in genes that evolved *de novo* and not in genes that diverged from ancestral coding sequences, we selected significant blast hit based on a relaxed threshold. Additionally, taking into account any discrepancies between gene annotations, we check for presence of *Triticeae* genes in the genomes of *Brachypodium* and rice. In order to exclude genes that were not annotated in the outgroup species, but that are actually present in their genomes, we mapped the CDS of *Triticeae* genes against the genomes of *Brachypodium* and rice using GMAP (version 2019-09-12). If ≥ 95% of the CDS, start codon and intact ORF were found in the genome of one of the two outgroup species, the gene was considered as “not *Triticeae* specific” and therefore filtered out.

Bi-directional Blastn analyses (query coverage 80%, identity 80%) between wheat, barley and rye, were used to identify genes that are homologous between at least two *Triticeae* species, and also to reduce the number of protein sequences to be analysed in the next step.

Finally, Blastp (evalue 1e-5) analyses of these *Triticeae* homologs against the non-redundant protein databases of NCBI and UniprotKb were performed to remove any proteins, that were not found in *Brachypodium* or rice, but that could be conserved outside of the *Triticeae* clade. Only genes with protein homology against members of the *Triticeae* clade or genes that do not show any sequence similarity with known proteins, were classified as candidate *de novo* TSGs and retained for further analyses.

### Transcriptomic analysis

Evidence of expression can be used to discriminate real genes from prediction artifacts. In addition, genes that are differentially expressed between stress conditions, are likely to be functional genes that are integrated into regulatory networks. Based on these assumptions, we measured the expression of the putative *de novo* TSGs under non-stress conditions and checked for differential expression under different abiotic and biotic stress conditions.

For quantifying the expression of *de novo* TSGs under non-stress conditions, we used gene expression values (transcripts per million, TPM; expression is normalized for gene length and sequencing depth) that were measured in the wheat lines Chinese Spring and Azhurnaya by Ramírez-González *et al*. (2018). The authors considered RNA-Seq data samples that were collected from 27 different tissues across 25 developmental stages (http://wheat-expression.com/download). First, we determined the average expression of each gene, by calculating the mean expression value across different developmental stages. This was done for each individual tissue. According to Ramírez-González *et al*. (2018), genes with an average expression value of >0.5 TPM in at least one tissue were considered as expressed. Secondly, we used these values for estimating the global expression of each gene (i.e., the average expression across all tissues in which a gene is expressed). Finally, we clustered genes in four different categories based on the following expression cut-offs: very weak expression (0.5-5 TPM), weak expression (5-10 TPM), average expression (10-20 TPM), and strong expression (>20 TPM).

Differential expression was measured against a broad range of abiotic and biotic stresses (Suppl. table S2). We considered wheat plants infected with major fungal pathogens, such as wheat powdery mildew (PRJNA296894, Praz *et al*., 2018; PRJNA243835, Zhang *et al*., 2014), *Zymospetoria tritici* (PRJNA327013, Ma *et al*., 2018; PRJEB8798, Rudd *et al*., 2015), *Fusarium graminareum (*PRJNA289545, *Gou et al*., *2016), Fusarium pseudograminareum* (PRJNA263755, Ma *et al*., 2014) and stripe rust (PRJNA243835, Zhang *et al*., 2014; PRJEB12497, Dobon *et al*., 2016). PAMP triggered immunity (PTI) response was measured from wheat plants treated with chitin and flagellin 22 (PRJEB23056, Ramírez-González *et al*., 2018), and finally cold (PRJNA253535, Li *et al*., 2015) was used to quantify abiotic stress responses.

*Triticeae* have large polyploid genomes and de novo genes are known to be associated with transposable elements (therefore several gene copies may be present at different loci within the genome). The software Salmon (Patro *et al*., 2017) is an alignment-free method which is known for its high computing efficiency and high accuracy of expression estimates for gene duplicates (Soneson *et al*., 2015) and isoforms (Sarantopoulou *et al*., 2021). Indeed, Salmon was shown to outperform “genome alignment-based” approaches (e.g., STAR+*featureCounts)* for the handling of multi-mapping reads (Soneson *et al*., 2015). For these reasons, we used Salmon for estimating the expression of wheat coding sequences (IWGSC RefSeq v1.0). Salmon was run in mapping-based mode with standard parameters and the estimated number of mapped reads (NumReads) were used for differential expression analysis as previously described by Praz *et al*. (2018). The analysis was performed with the R package edgeR (Robinson et al., 2009). The calcNormFactors function was first used to normalize the estimated number of mapped reads to TMM values (Trimmed Mean of M-values). Only genes with >5 CPM (count per millions) in at least 3 RNA-Seq samples were retained. We used the functions estimateGLMCommonDisp, estimateGLMTrendedDisp, and estimateGLMTagwiseDisp for estimating dispersion, and fitted a negative binomial generalized linear model with the glmFit function. Finally, likelihood ratio test (glmLRT function) was used for determining differential gene expression through different pairwise comparisons. Only genes with a log2FC > |1.5| and an adjusted *p*-value (FDR) < 0.01 were considered as differentially expressed.

Finally, we plot a heatmap to compare the response of differentially expressed TSGs across different stress treatments (Fig. 2a). For each gene, we considered the log2FC value that was calculated between test and control conditions, and the R package pheatmap was used for generating the plot. In order to improve the data visualization, log2FC values were scaled to Z-scores with the argument scale = “row”.

### dN/dS and translation evidence

Evidence of translation and/or negative selection pressure are strong indications that a gene is functional and encodes a protein. The codeml program of the PAML package (Yang, 2007) was used for measuring the interspecies selection pressure between homologous TSGs of rye, barley and wheat. The strength of selection pressure was measured as the ratio between nonsynonymous (dN) and synonymous amino acid changes (dS). For selecting TSGs that are under negative selection pressure, we considered the median dN/dS value of all TSGs that are supported by proteomic evidence (i.e., TSGs that are under evolutionary constraints; Suppl. fig. S2). The median of the distribution was found at ∼0.53, therefore all TSGs that fall below this value were considered as genuine protein-coding genes. Finally, protein sequences of TSGs were aligned (Blastp, evalue 1e-10) against the wheat proteomic database (UniProt, UP000019116).

### *In silico* characterization of TSGs

*De novo* genes originate from non-coding sequences and therefore is not possible to infer its function based on sequence homology with previously described genes. In order to investigate the putative function of the identified TSGs, we predicted the presence of conserved domains. The algorithms SignalP (v5.0) and TMHMM (v2.0) were used to annotate signal peptide (SP) and transmembrane (TM) domain. Given that N-terminal TM domains can be misannotated as SP, we manually curated TSGs that were predicted to contain both domains. More specifically, genes in which i) at least one TM and no SP are predicted, ii) one SP and more TMs are predicted, and ii) the SP have a low probability score (<0.5), were considered as genes encoding transmembrane proteins. Whereas genes in which i) one SP and no TMs, or ii) one SP and one N-terminal TM domain are predicted, were considered as apoplastic proteins (Suppl. fig. S3).

Orthofinder (v2.3.12; Emms & Kelly, 2015) was used for clustering groups of orthologous TSGs (i.e., orthogroups, hereafter called gene families), based on protein sequence homology. Orthofinder was run with standard parameters using the protein sequences (one file per *Triticeae* species) of candidate TSGs. To identify *de novo* genes that might have originated from TEs, we aligned nucleotide and protein sequences of TSGs against consensus TE sequences from the nrTREP database and their derived proteins PTREP (botinst.uzh.ch/en/research/genetics/thomasWicker/trep-db.html). TSGs that have homology (Blastn, identity > 80%) at the nucleotide level, but did not show any similarity (Blastp, evalue 1e-10) with TE proteins, are likely to have originated from TE non-coding regions (e.g., LTRs and TIRs) or through frame-shift mutations in TE coding regions. In order to distinguish them, we further aligned (Blastx, evalue 1e-10) nucleotide sequences of these TSGs against PTREP. Genes with significant Blastx homology against canonical TE proteins likely evolved through frame-shift mutations and were classified as “out-of-frame” TSGs (Table 2).

## Results

We defined *Triticeae*-specific genes (TSGs) as genes that are annotated in at least two of the main *Triticeae* lineages represented by wheat, rye and barley, but have no homologs outside of the *Triticeae* clade, in particular in the closely related outgroup grasses *Brachypodium distachyon* and rice (Suppl. fig. S1). In this study, we focused exclusively on protein coding genes and excluded non-coding RNAs or small RNAs. For our analyses we used the recently published chromosome-scale genome assemblies of bread wheat, barley and rye (IWGSC *et al*., 2018; Mascher et al., 2017; Rabanus-Wallace *et al*., 2021). As outgroups we used *B. distachion*, rice, and the plant protein databases of NCBI and UniprotKb. First, we assessed the integrity of the coding sequence (CDS) prediction, by examining whether they indeed represent intact open reading frames (ORFs). In wheat low-confidence genes, approximately 10% contained in-frame stop codons (Suppl. table S1), suggesting the predicted CDS was in the wrong reading frame. These were discarded. Depending on the species, between 2% (wheat) and 44% (barley) of the predicted CDS lacked start or stop codons (Suppl. table S1). Genes missing stop codons were still used for the analysis, the others were discarded. Discrepancies between *Triticeae* gene annotation strategies (see methods) might explain why barley has the largest number of genes with incomplete ORF. To exclude all genes that might have originated in ancestral grasses, we ran sequence similarity searches against the genomes (GMAP, see methods) and the annotated protein sequences (Blastp, e-value 1<10E-5) of both outgroup species *Brachypodium* and rice. *Triticeae* genes that have sequence homology in at least one of the outgroup species were eliminated. This resulted in a data set of 67,413 candidate TSGs. As expected, the majority (61,508or ∼91%) of these were previously annotated as low confidence genes.

As a first selection for candidate TSGs, we performed all vs. all bi-directional Blastn searches between the CDS of wheat, rye and barley and identified the best homologous gene pairs between the three genomes. Here we only kept genes that are annotated in at least two of the three genomes. Finally, we ran an additional Blastp analysis between the identified gene homologs and the protein databases of NCBI and UniprotKb and discarded all genes that had homologs conserved outside of *Triticeae* tribe. In total, we found 5,942 genes that are commonly predicted in at least two of the *Triticeae* species, but not in other plant species outside of the *Triticeae* tribe. This initial dataset of candidate TSGs comprised 3,680 genes from wheat, 1,819 from rye 443 from barley (Table 1, Fig. 1).

**Table 1.**
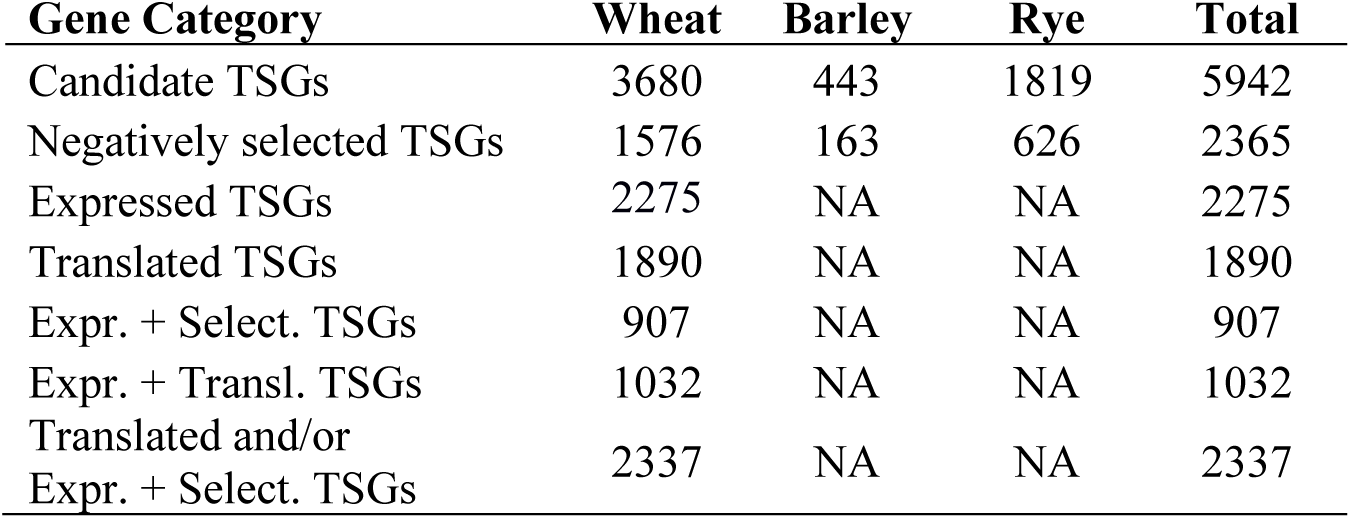
Summary of predicted and *in silico* characterized *Triticeae*-specific genes (TSGs). Candidate TSGs were identified by selecting all annotated genes that are conserved in at least two *Triticeae* species (wheat, barley and rye), but are not found in other plant species outside of the *Triticeae* clade (see methods). Candidate TSGs were further characterized, by measuring 1) gene expression in wheat under non-stress conditions, 2) sequence homology to the wheat proteomic database (translated TSGs) and 3) evidence of negative selection. Candidate TSGs with experimental evidence of translation and/or evidence of both transcription and negative selection, were considered as “real” protein-coding genes. Therefore, we suggest that *Trititiceae* may possess ∼779 (2,337 / 3 subgenomes) lineage-specific genes per haploid genome.

**Fig. 1.**
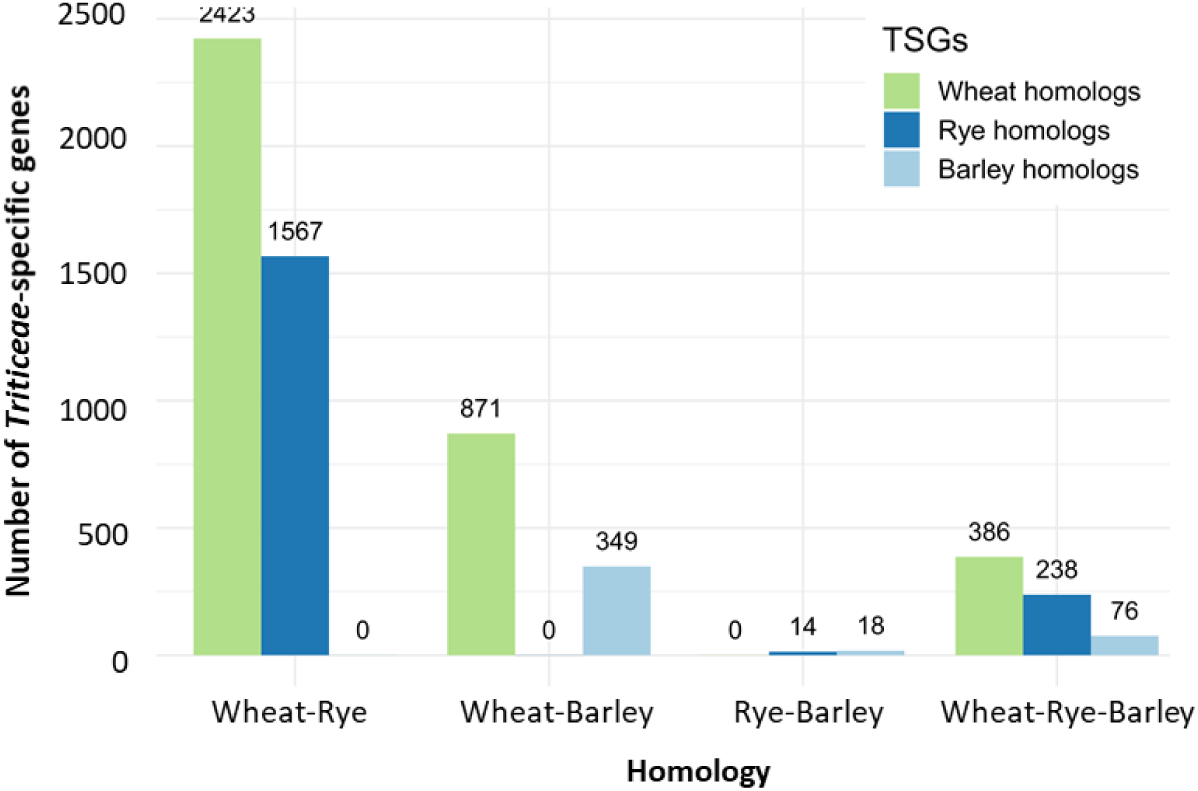
Candidate *Triticeae*-specific genes (TSGs) identified between wheat, barley and rye. Bi-directional Blastn of genes that are not conserved outside of the *Triticeae* clade, resulted in a total of 3,680, 1,819, and 443 genes corresponding to wheat, rye and barley, respectively. TSGs were defined as genes which have homology in at least two *Triticeae* species but are not conserved outside of the *Triticeae* clade. Therefore, the bars show the total number of TSGs per species which are shared between *Triticeae*. Between wheat and rye, for example, 2,423 wheat genes found homology with 1,567 rye genes, and none those were annotated in the barley genome. Note that only a small proportion of genes (700 TSGs) were commonly found in all three *Triticeae* species.

Homology relationships at the protein level between candidate TSGs, were defined using OrthoFinder (Emms & Kelly, 2015). In total, predicted proteins of 2,555 TSGs were clustered into 857 orthogroups (hereafter families) with at least two members each. In the remaining 3,387 unassigned TSGs, no homology at the protein level was observed.

### Many candidate TSGs are supported by dN/dS analysis, expression and proteomic data

Since real protein-coding *de novo* genes should be expressed, translated and under purifying selection (McLysaght & Guerzoni, 2015; Prabh & Rödelsperger, 2016) we searched for support in transcriptomic and proteomic data, and calculated the strength of selection between homologs from the three species. For the transcriptomic analysis, we used published expression data (transcript per million TPM values) that was collected in two different wheat lines Azhurnaya and Chinese Spring from a total of 27 tissues and 25 developmental stages (Ramírez-González *et al*., 2018). Given that many functionally characterized genes are low expressed under normal conditions, the threshold of minimum expression was set at “> 0.5 TPM in at least one tissue” as previously proposed by Ramírez-González *et al*. (2018). Using this threshold, 2,275 out of 3,680 wheat TSGs (∼62%) showed expression, with 1,774 being very weakly (0.5-5 TPM), 1,976 weakly (5-10 TPM), 306 average (10-20 TPM), and 266 highly expressed (> 20 TPM) in at least one of the two wheat lines. In a second step, we searched for matching protein sequences in the available proteome database of wheat (UP000019116). In total, we found that 1,890 out of 3,680 wheat TSGs (∼51%) have strong matches in the proteomic data. Of these, 1,032 wheat TSGs also show transcriptomic evidence in the RNA-Seq data (Table 1).

Additionally, we used pairwise comparisons between TSGs from wheat, barley and rye homologs to determine the ratio of substitutions at non-synonymous and synonymous sites (dN/dS = ω). By choosing a stringent cut-off of ω = 0.53 (see methods and Suppl. fig. S2), we found that 2,365 out of 5,942 TSGs (∼40%) are under evolutionary constraints; 1,576 TSGs are from wheat, of which 907 (∼58%) also show evidence of expression.

Experimental evidence of translation is the strongest indication that a gene is real, and it can be used to validate the protein-coding potential of genes. In addition, according to Prabh & Rödelsperger (2016), we also considered TSGs that show both evidence of expression and negative selection (see above) to be real protein-coding genes. Indeed, a large proportion (460 out of 907; 51%) of these “expressed and negatively selected” genes is also supported by proteomic evidence. To summarize, ∼64% (2,337 out of 3,680) of the candidate wheat TSGs were translated and/or supported by both evidence of expression and negative selection. These candidate TSGs were therefore validated as genuine protein-coding genes (Table 1). We carefully propose that *Trititiceae* may possess ∼779 (2,337 genes / 3 subgenomes) lineage-specific genes per haploid genome.

### Stress-induced TSGs are enriched in secreted proteins

Signal peptides (SP) are a subclass of N-terminal transmembrane (TM) domains which mediate the secretion of proteins into the apoplast. In contrast to TM domains, SPs are cleaved off after translocation through the ER membrane. Mature proteins can ultimately be released outside of the cellular membrane through the classical secretory pathway (Wang *et al*., 2018). However, because of structural similarities between SPs and TM helices, these two N-terminal signaling sequences are often miss-annotated or annotated inter-changeably (Yuan *et al*., 2003). Indeed, we found that 286 candidate TSGs were ambiguously predicted to encode both N-terminal TM domain and SP (Suppl. fig S3). Manual curation (see methods and suppl. fig S3) revealed that ∼37% (2,204 out of 5,942) of our candidate TSGs might encode secretory proteins. Of these, 1,622 and 582 are likely to be located at the plasma membrane or in the apoplast, respectively. Interestingly, we found that 14 of the 35 most abundant TSG families (≥ 10 gene members), including the three largest ones, encode putative transmembrane proteins (Suppl. table S5). This suggests that the presence of transmembrane domains might have played an important role in the evolution and diversification of *de novo* genes.

Differential gene expression analyses can be used to define genes which are likely to be embedded into regulatory networks and therefore to indicate their function (Prabh & Rödelsperger, 2016). Because of the large availability of RNA-Seq datasets for wheat, we focused on the 3,680 candidate TSGs from wheat. The publicly available datasets (Praz *et al*., 2018; Ma *et al*., 2018; Ramírez-González *et al*., 2018) include wheat exposed to cold stress, to pathogen associated molecular patterns (PAMP; chitin and flg22) elicitation as well as wheat that was infected with different fungal pathogens (see methods and Suppl. table S2).

In total, we found 215 TSGs that were significantly differentially expressed (Log2FC > 1.5, FDR < 0.01) in at least one stress treatment, 133 TSGs were up-regulated and 82 were down-regulated. DE genes that belong to the same families, show similar patterns of transcriptomic response, suggesting some degree of functional redundancy within TSG families (Fig. 2A, Suppl. table S3). As we were interested in TSGs that are specifically induced upon stress, we further characterized 133 genes that were up-regulated in at least one treatment. To our surprise, stress-induced TSGs are enriched in secreted proteins, as ∼50% of them are predicted to be secreted to the apoplast (35 TSGs) or to localize to the cell membrane (32 TSGs). Most interestingly, specific patterns of gene expression seem to distinguish these two groups: most putative transmembrane proteins are up-regulated by biotic stresses such as *Z. tritici* infection and PAMP (chitin and flg22) treatment, whereas apoplastic proteins are mainly induced by cold stress and stripe rust infection (Fig. 2A-B).

**Fig. 2.**
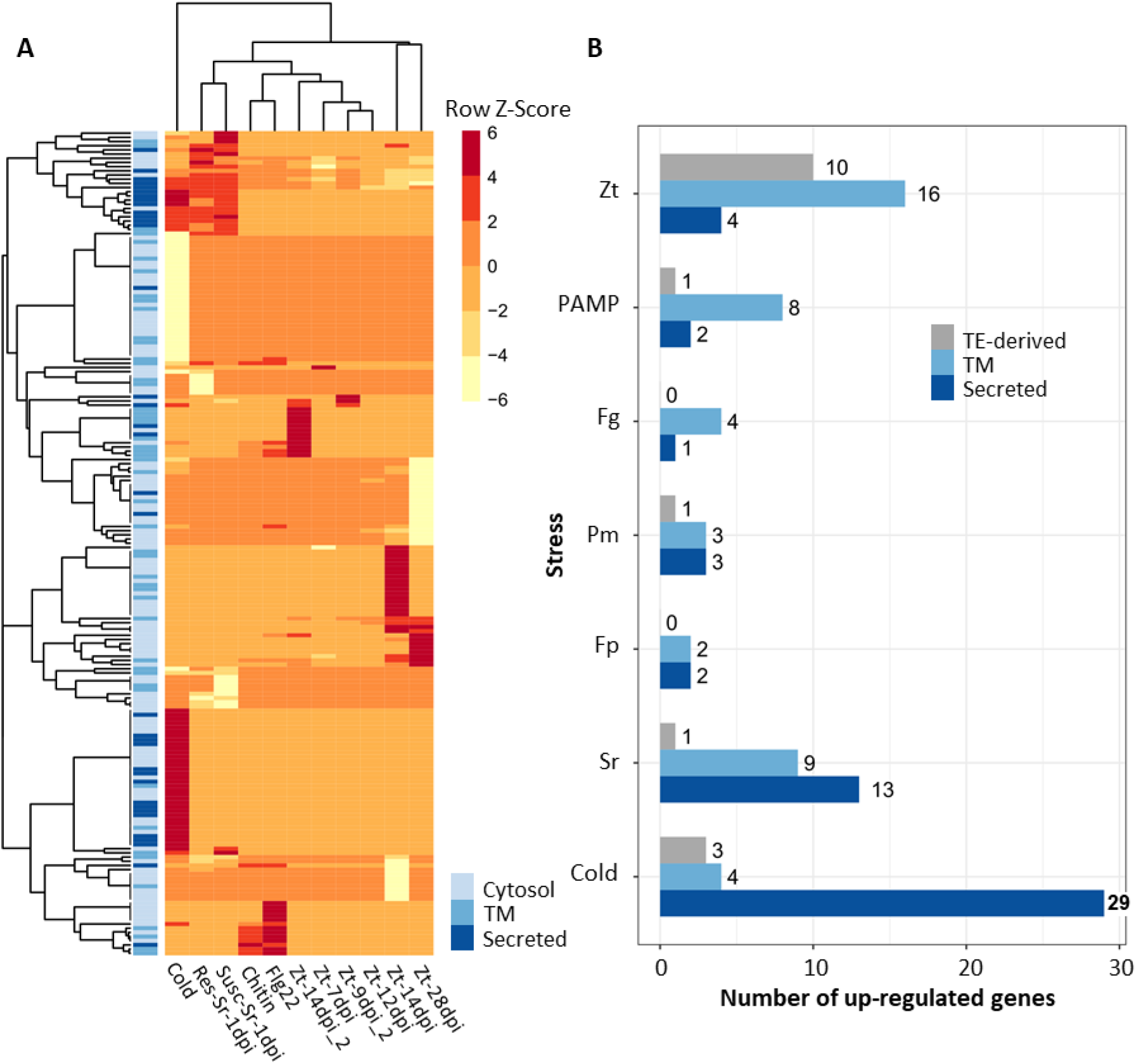
Transcriptomic analysis of differentially expressed *Triticeae*-specific genes (TSGs). **(A)** Heatmap showing the transcriptomic response of differentially expressed (DE) TSGs. The R package edgeR was used for the DE analysis of candidate TSGs. Only genes that were significantly up-regulated (log2FC > |1.5| and adjusted *p*-value FDR < 0.01) and only the treatments that showed high transcriptomic response, were visualized in the heatmap. The R function pheatmap was used to scale the log2FC values into Z-scores and to draw the plot. In addition, DE genes were annotated based on on the predicted cellular localization (transmembrane or secreted apoplastic proteins). **(B)** Bar plot showing the number of secreted, transmembrane (TM) and TE-derived genes that are up-regulated upon stress. Zt: *Z. tritici*; PAMP: pathogen associated molecular patterns; Fg: *F. graminareum*; Fp: *F. pseudograminareum*; Pm: powdery mildew; Sr: stripe rust.

### TSGs are involved in climate adaptation and non-host resistance of *Triticeae* species

Interestingly, 23 out of 29 apoplastic TSGs that are induced by cold stress, belong to the “low-temperature or cold-responsive” *Lt-Cor* gene family (Fig. 3A). *Lt-Cor* genes are cereal-specific glycine-rich proteins which are known for increasing freezing tolerance and therefore being involved in the low-temperature acclimation of cereals (Ohno et al., 2001; Pearce et al., 1998). Based on sequence homology we clustered these genes in three subgroups (Fig. 3A): 8 wheat TSGs that show strong similarity to barley and rye members of the *lt14* gene family (Dunn *et al*., 1990), 2 wheat TSGs that contain a stretch of contiguous repeats of a “LPT” triplets which is characteristic of the wheat homolog *Wlt10* (Ohno et al., 2001), and 13 wheat TSGs that belong to the *Tacr7* family (Gana *et al*., 1997). Furthemore, we found that 12 of these genes (many of which belonging to the *Tacr7* family) show up-regulation after both cold stress and stripe rust infection (particularly at 1 dpi; Fig. 2A). Stripe rust is specialized to infect wheat at cool temperatures (Gaudet *et al*., 2011), therefore further suggesting that TSGs of the *Lt-Cor* gene family are involved in stress responses against cold and cold-adapted fungal pathogens.

**Fig. 3.**
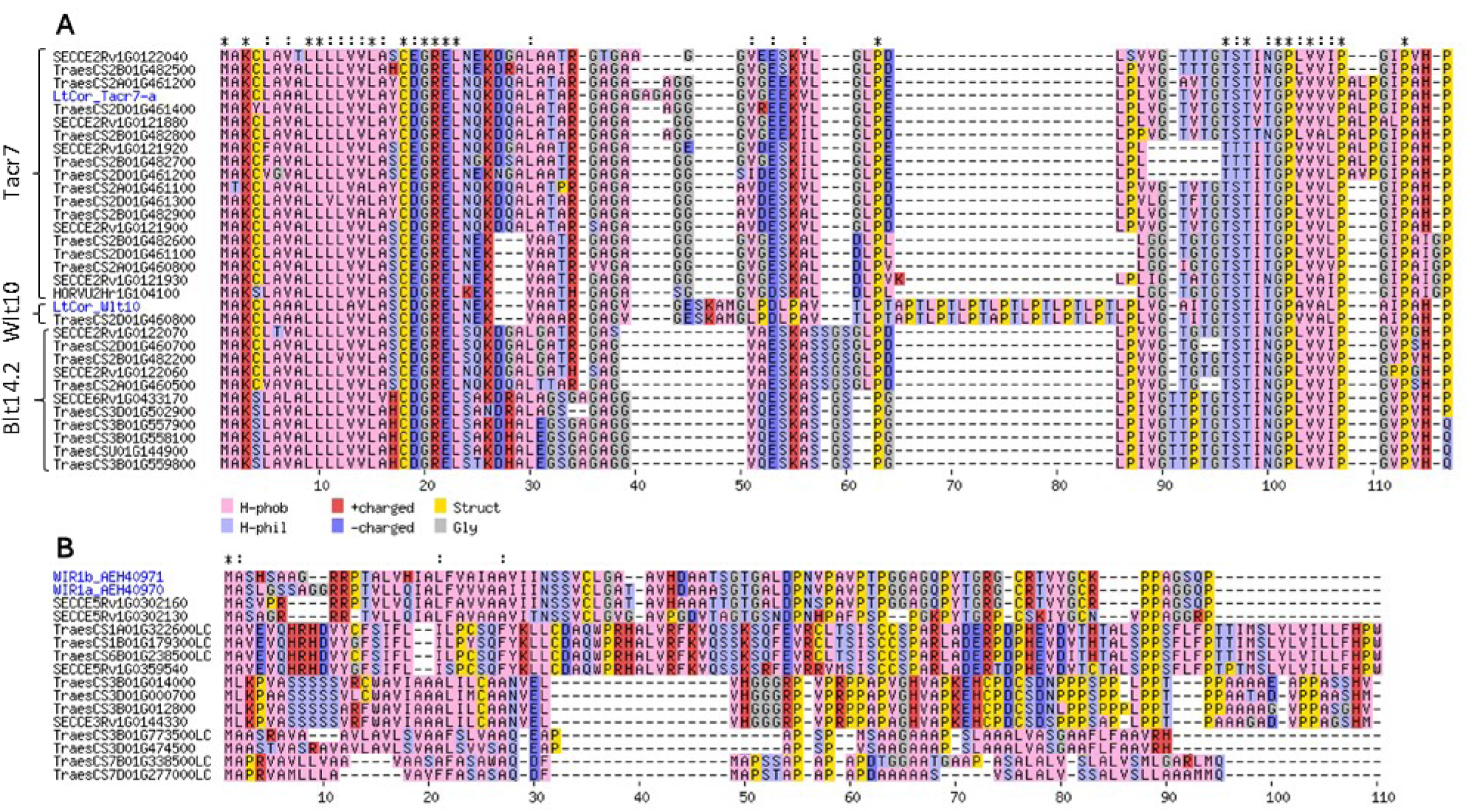
Multiple alignment of predicted protein sequences of known stress-induced *Triticeae*-specific gene (TSG) families. **(A)** “Low-temperature or cold-responsive” *Lt-Cor* gene family; previously characterized proteins (TraesCS2A01G460600: Wlt10, AAF75555.1; TraesCS2B01G483000: Tacr7-a allele, AWT24553.1) are highlighted in blue. **(B) “**Wheat-Induced Resistance 1” WIR1-like proteins; previously characterized WIR1 proteins (TraesCS5B01G045100: TaWIR1a, AEH40970.1; TraesCS5D01G050200: TaWIR1b, AEH40971.1) are highlighted in blue.

In contrast to “apoplastic” TSGs, membrane-localized TSGs encode a set of very diverse proteins which were classified into 24 different gene families. In particular two membrane-localized TSGs (TraesCS5B01G045100, TraesCS5D01G050200) stood out for showing the strongest stress response: they are up-regulated in response to several hemibiotrophic and biotrophic fungal pathogens (*Fusarium*, powdery mildew and stripe rust) and also after exposure to PAMPs (chitin and flg22). These genes encode proteins with strong homology to the two WIR1 (Wheat-Induced Resistance 1) proteins TaWIR1a (GenBank accession AEH40970.1) and TaWIR1b (GenBank accession AEH40971.1; Fig. 3B). The *WIR1* gene family is known to confer non-host resistance against a wide range of fungal pathogens, and it was suggested to be involved in basal resistance of cereals (Tufan *et al*., 2012). In addition to the already described wheat (*TaWIR1a/b*; Tufan et al. 2012) and barley (*HvWIR1*; Douchkov *et al*., 2011) homologs, we also identified two novel *WIR1* genes in rye (SECCE5Rv1G0302160 and SECCE5Rv1G0302130, Fig. 3B). Furthermore, we identified 12 novel WIR1-like proteins which contain transmembrane domains, have similar Proline-and Glycine-rich motifs, and are also induced after *Z. tritici* and PAMP treatments (Fig. 3B).

### Several TSGs are derived from non-coding regions of transposable elements

Because previous studies suggested transposable elements (TEs) to be involved in many ways in the evolution of lineage-specific genes (Toll-Riera *et al*., 2009; Tautz & Domazet-Lošo, 2011; McLysaght & Guerzoni, 2015; Jin *et al*., 2019), we were particularly interested in TSGs deriving from such repetitive elements. We found that approximately 18% of TSGs (1,079 out of 5,942) show nucleotide sequence homology to TEs (Table 2). Interestingly, the majority of these genes (1,067 out of 1,079; 99%) do not encode any canonical TE proteins (Table 2). More than ∼95% of TGS in barley (119 out 121) and wheat (841 out of 880) are located in non-coding portions of TEs. In rye ∼46% (36 out of 78) are predicted in alternative frames of canonical TE proteins and ∼51% (40 out of 78) are found in the non-coding regions of TEs. Taken together, these data suggest that sequence divergence from non-coding regions and frame-shift mutations in existing ORFs (particularly in rye) might be an important source of novel gene evolution from *Triticeae* TEs.

**Table 2.** Number of *Triticeae*-specific genes (TSGs) that are associated to transposable elements (TEs). Here we summarize the number of TSGs that show homology at the DNA and protein level with TEs. The majority are found in non-coding regions (e.g., TIRs and LTRs) or are out-of-frame with canonical TE proteins (transposases, integrases, etc.). In addition, we indicate the number of TSGs which are predicted to encode at least one transmembrane domain (TM).

|  | Wheat | Barley | Rye | Total |
| --- | --- | --- | --- | --- |
| DNA homology | 880 | 121 | 78 | 1079 |
| Protein homology | 9 | 1 | 2 | 12 |
| “Out-of-frame” TSGs | 30 | 1 | 36 | 67 |
| “Out-of-frame” TSGs - TM | 6 | 0 | 29 | 35 |
| “Non-coding regions” TSGs | 841 | 119 | 40 | 1000 |
| “Non-coding regions” TSGs - TM | 180 | 30 | 6 | 216 |

Out of 1,079 TE-derived TSGs, we identified 25 differentially expressed genes. In total, 9 of them were down-regulated, whereas 12 were induced by fungal pathogens, 3 by cold stress and 1 by PAMP treatment (Fig. 2B). Despite the low number, we found that 37.5% (6 out of 16) of these up-regulated TSGs encode TM domains and are all induced by *Z. tritici* infection (Fig. 2B). This further suggests that TM-encoding TSGs are particularly involved in biotic stress responses. The up-regulated TE-derived genes were analysed with special scrutiny, through detailed annotation of the regions containing the predicted genes and analysis of expression data. These genes are members of 8 different families (TE1 through TE8), that include a total of 107 TSGs. The gene families have between 2 (TE3) and 57 members (TE7). Detailed analysis was performed of those family members that show differential expression. Gene families TE1 through TE4 encode proteins with predicted transmembrane domains, while T5 though TE8 encode no identifiable protein domains.

For gene family TE2, we have the most complete understanding of its evolution. TE2 is a gene family of 15 genes in wheat and 2 in barley. All genes are predicted upstream of the transposase gene in *CACTA* transposons of the *DTC_Clifford* family (Fig. 4A), eight are expressed under non-stress conditions in wheat, and three are induced upon infection with *Zymoseptoria* and one after PAMP elicitation. The example of TraesCS7A01G416500 shows that expression is limited to the region of the predicted gene plus its upstream region, while the rest of the *DTC_Clifford* element is transcriptionally silent (Fig. 4A). Our data indicate that the existence of this gene family is the result of a sequence capture event. We propose that the *DTC_Clifford* family captured two probably non-coding sequences and fused them to a TE2 family proto-gene. This presumably occurred in the *Triticeae* ancestor, because *DTC_Clifford* transposons from wheat and barley all contain that segment, while in contrast, the closest *DTC_Clifford* homologs in oat do not (Fig. 4C). Interestingly, we found the putative progenitor sequences in the oat genome: the 5’half of the proto-gene is present in two copies on chromosomes 6C and 7D, while homologs of the 3’ half are found in five copies on chromosomes 1A, 3A, 4A, 5A and 5D. The captured sequences also contain a lower G/C content than the surrounding sequences (Fig. 4B). The newly captured and fused sequence was then amplified along with the *DTC_Clifford* elements. From these ancestor sequence, putatively functional TE2 family genes evolved both in wheat and barley.

**Fig. 4.**
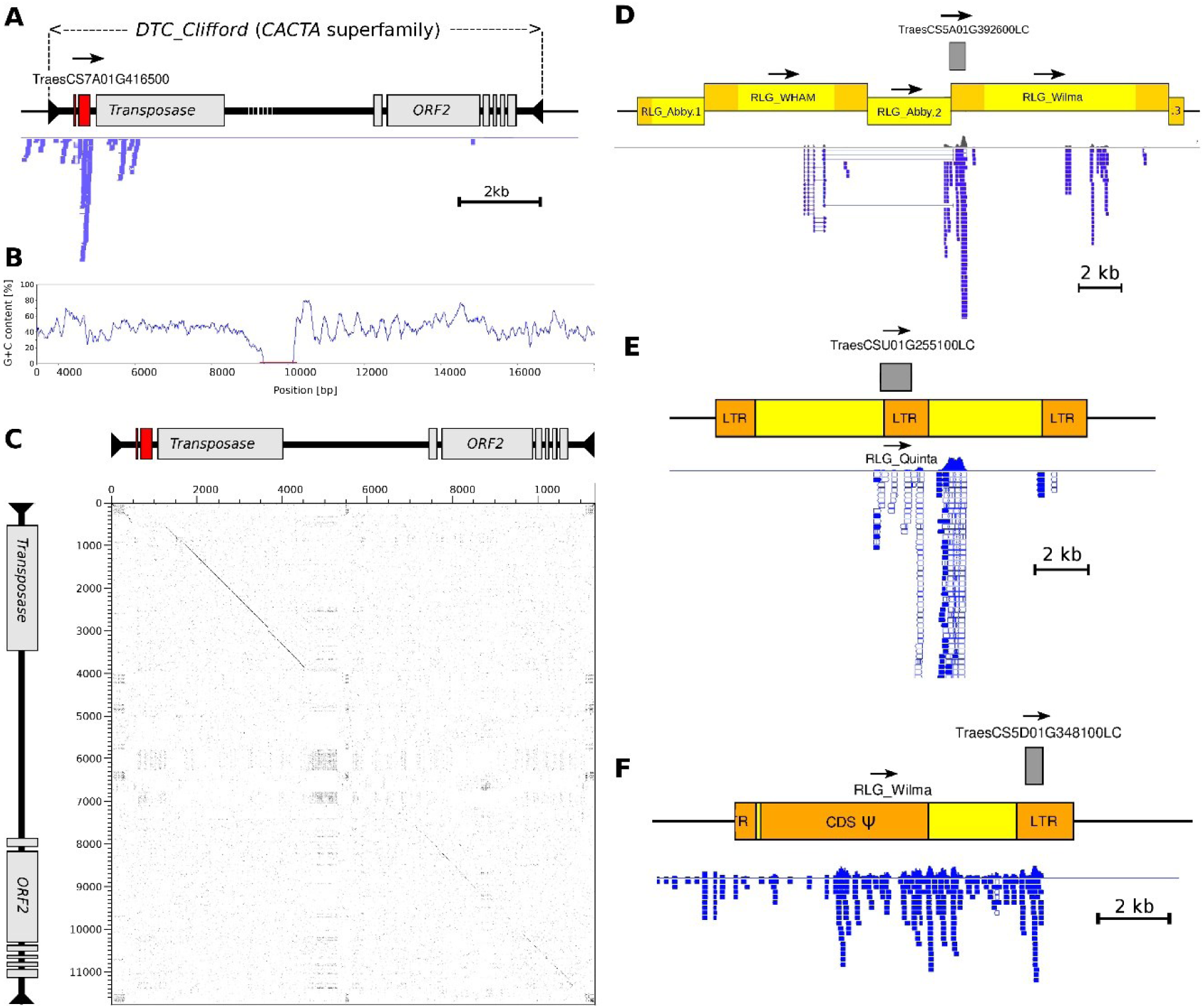
Genomic and transcriptomic analysis of TE-derived *de novo* genes which are induced upon infection with *Zymoseptoria*. **(A)** Sequence organization of TE2 gene family members. The predicted gene *TraesCS7A01G416500* (exons as red boxes) is located just upstream of the CDS for transposase inside *CACTA* transposons of the *DTC_Clifford* family. Mapped transcriptome reads are shown in blue underneath the map, confirming the predicted exon/intron structure. **(B)** Plot of G/C content in 100 bp windows with a 10 bp sliding step. Note that the region of the predicted gene has a lower gene content that the surrounding sequence. **(C)** Dot plot comparison between *DTC_Clifford* from wheat (horizontal) and its closest homolog from oat (vertical). The region containing the predicted gene is absent from the oat homolog, indicating that the sequence was captured by *DTC_Clifford* in the *Triticeae* lineage. **(D-F)** Examples of differentially expressed genes located in LTRs or retrotransposons. The position of the predicted gene is indicated with a grey box above the map, and mapped transcriptome reads are shown in blue underneath the map. Despite the overall high copy number of the retrotransposons, unique sequence combinations can arise through nested insertions (**D**), duplications (**E**) or deletions that truncate elements **(F)**.

Families TE1, TE5 and TE7 are derived LTR sequences of *Gypsy* retrotransposons (Fig. 4D-F). Gene predictions in such highly abundant sequences are notoriously suspicious. Nevertheless, at least some copies have specific and unique characteristics: gene TraesCS5A01G392600LC is predicted across the boundary of two LTR retrotransposons, where an *RLG_Wilma* retrotransposon inserted into an *RLG_Abby* retrotransposon (Fig. 4D). Because retrotransposons insert mostly randomly, such TE-junctions are highly specific. Interestingly, RNA-seq data shows gene expression across that junction and only for the region of the predicted gene while most of the surrounding TE sequences are transcriptionally silent (Fig. 4D). Similarly, TraesCSU01G255100LC is located in an *RLG_Quinta* tandem repeat (i.e., a recombined element with 3 LTRs and 2 internal domains; Fig. 4E). Expression is only found in the region of the predicted gene as well as the *gag* region of one internal domain, but not in the rest of recombined elements (Fig. 4E). These data suggest that these genes are indeed functional *de novo* genes.

### Transmembrane domains are derived from repetitive AT-rich regions

The majority (1,000 out 1,079; ∼93%) of TE-derived TSGs were found inside or close to regulatory regions such as long terminal repeats (LTRs) and terminal inverted repeats (TIRs, Table 2). LTRs and TIRs are known to contain short but highly conserved sequences that are important for the transcription and mobilization of TEs (e.g., AT-rich regions at the termini of LTRs and TIRs, and poly-adenylation signals; Benachenhou et al., 2013; Wicker et al., 2007). Of these 1,000 genes from non-coding regions of TEs (961 being LC), we found that 216 (21.6%) are predicted to encode at least one transmembrane domain. In contrast, only 9.5% of the overall wheat LC genes encode predicted transmembrane domains (Table 2).

Interestingly, these TM-encoding genes have unusually low GC contents (Fig. 5C-D). Although LC genes have a generally lower GC content than HC genes, the TE-derived genes (which mostly are LC genes) are among those with the lowest values (Fig. 5B). Similarly, the TM-encoding TE-derived genes have much lower GC contents than TM-encoding HC and LC genes (Fig. 5C). Indeed, AT-rich sequences are generally likely to give rise to ORF encoding transmembrane domains, as the hydrophobic amino acids are mostly encoded by codons with at least 2 Adenines or Thymines (Prilusky & Bibi, 2009; Vakirlis *et al*., 2020). Since non-coding regions of TEs tend to be AT-rich, we propose that novel ORFs evolving from them are more likely to encode a transmembrane domain.

**Fig. 5.**
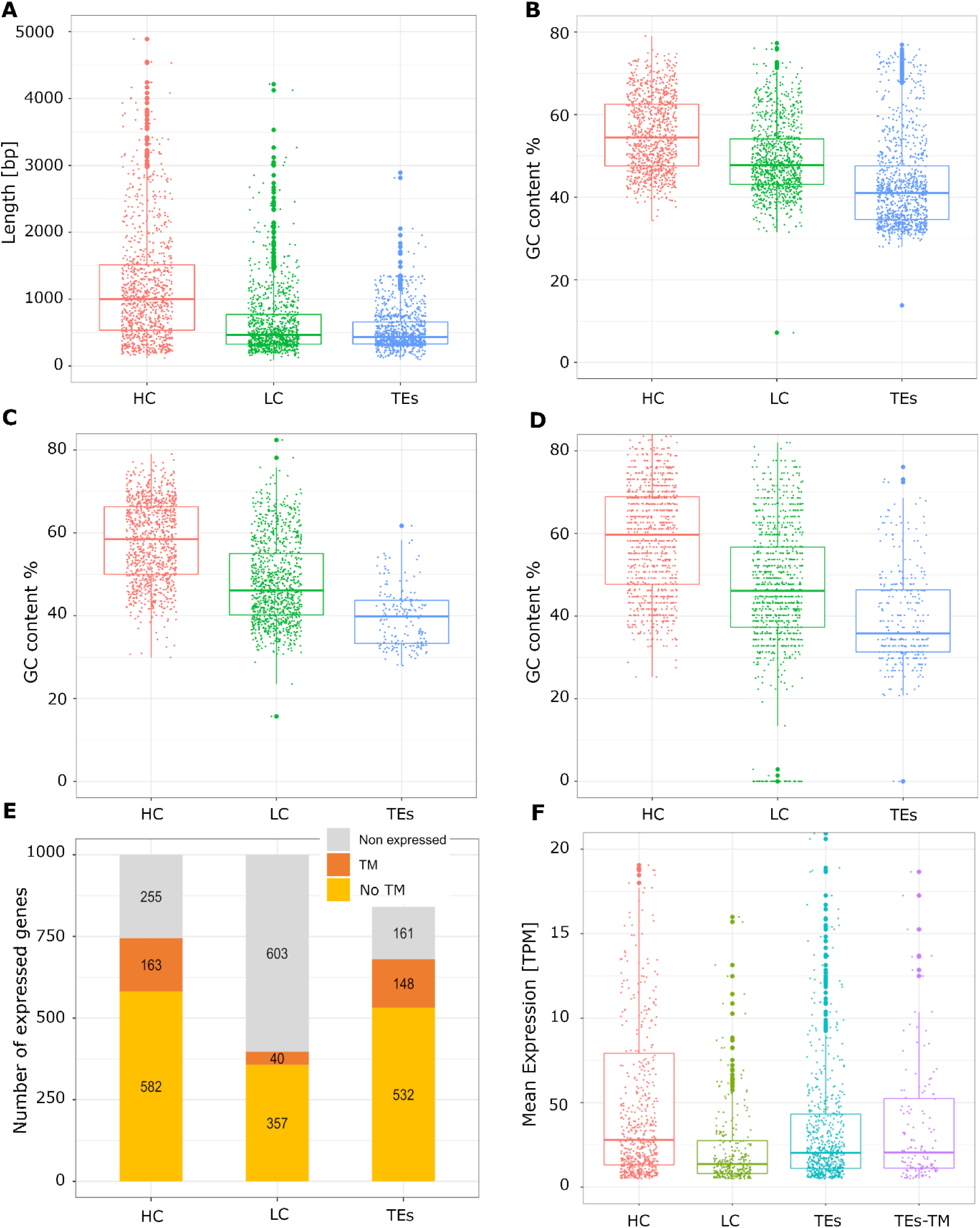
Transcriptomic and GC-content analyses of TSGs coming from TE non-coding regions. Comparison between 1,000 *Triticeae*-specific genes (TSGs) derived from transposable elements (TEs) non-coding regions and 1,000 randomly selected low-confidence (LC) and high-confidence (HC) genes. **(A)** Size comparison, **(B)** overall GC-content, **(C)** GC-content of genes that encode transmembrane (TM) domains **(D)** GC-content from the region encoding the TM domains. **(E)** Number of expressed genes in wheat. Orange: TM-encoding genes; yellow: non TM-encoding genes; grey: remaining non-expressed genes. **(F)** Average expression value in the wheat line *Chinese spring* (see methods). TEs-TM: TE-derived genes encoding TMs.

Moreover, we observed that wheat TSGs from TEs non-coding regions are strongly supported by transcriptomic data. Approximately 81% (680 out of the 841) of them show expression in at least one wheat line (Fig. 5E), and their average expression level is higher than in most LC genes and it resembles that of HC genes (Fig. 5F). The overall fraction of expressed TE-derived genes is much higher than in LC genes (397 out of 1,000 randomly-selected LC genes, ∼40%) and even higher than in HC genes (745 out of 1,000 randomly-selected HC genes, ∼75%; Fig. 5E). Surprisingly, some of these genes, especially the ones encoding TM domains, showed among the highest expression levels of all wheat genes (Fig. 5F).

To summarize, our results indicate that the vast majority of TE-derived TSGs are located in non-coding regions of TEs. These genes are enriched in TM-encoding domains and are strongly supported by transcriptomic data, therefore supporting their functionality and protein-coding potential. For these reasons, we suggest that TEs non-coding regions might promote the origination of functional *de novo* genes by providing genomic raw material from which TM-encoding ORFs can evolve.

## Discussion

*Triticeae* form an ideal model system to study the evolution of *de novo* genes for several reasons: (i) they are closely enough related so that direct comparison of gene content and collinearity is possible, but yet distant enough to have numerous differences, (ii) chromosome scale genome assemblies and genome annotations of closely related outgroup species (*B. distachyon* and rice) are available, and (iii) up to 85% of the genome consists of transposable elements (TE) which were shown to be main drivers of genomic diversity in *Triticeae* (Wicker et al., 2017, 2018). Furthermore, as all crop plants, *Triticeae* are under evolutionary pressure to adapt to pathogens and changing environmental conditions.

Lineage-specific *de novo* genes have by definition no homologs outside of their respective taxonomic group. Thus, the most challenging aspect is to distinguish potential real genes from annotation artifacts. We therefore defined a set of stringent criteria to identify candidate *de novo* genes. Our first filtering step was to only consider genes with intact open reading frames, and whose nucleotide and protein sequences have no homologs in plant species outside of the *Triticeae* clade, in particular in the closely related *B. distachion* and rice. Using nucleotide sequence homology between *Triticeae* species (wheat, barley and rye) as an additional criterion, eliminated over 90% of the annotated low-confidence genes and resulted in the final set of 5,942 candidate TSGs. We used additional filters such as signatures of purifying selection, and evidence for transcription and translation to further measure the protein-coding potential of these candidate TSGs. Our approach differed form that in a recent study (Ma *et al*.,2020) which identified 3,812 candidate TSGs using expressed sequence tags (ESTs), but did not further verify for their protein-coding potential. ESTs do not cover the transcriptome as well as RNA-seq data (Ping *et al*., 2012; Lowe *et al*., 2017), especially for lineage-specific genes, which are often expressed at low levels or only under specific conditions. Furthermore, in contrast to Ma *et al*. (2020), we considered it crucial to include TE-derived TSGs, because studies in primates and rice have shown that over 50% of lineage-specific genes have homology to TEs (Toll-Riera *et al*., 2009; Jin *et al*., 2019). Indeed, we found that ∼18% of our identified TSGs are derived from TEs (mostly from non-coding regions), indicating that also in *Triticeae* TEs might be an important contributor to *de novo* gene evolution. Finally, we validated the protein-coding potential of most wheat TSGs, and we cautiously propose that *Triticeae* may contain roughly 779 lineage-specific genes per haploid genome.

It is very difficult to predict the function of *de novo* genes due to lack of sequence homology to functionally described genes. Because we were interested in the identification of TSGs that might be involved in responses to diseases or adaptation to environmental conditions, we used RNA-Seq data from wheat plants that were either infected with fungal pathogens, artificially treated with PAMPs or exposed to cold temperature. Interestingly, we found that up-regulated TSGs are enriched in genes encoding proteins with signal peptides and transmembrane domains, suggesting them to be transmembrane or apoplastic proteins. The apoplast is the interface between the plant cell membrane and many microbial parasites, and it is also the place where most freezing related responses take place (Kuwabara & Imai, 2009). Thus, it makes sense for novel genes encoding secreted and transmembrane proteins to be functionally selected during stress responses. In fact, similar results were found in yeast where adaptive (fitness-enhancing) sequences were enriched in TM domains, leading to the development of the “TM-first” model of *de novo* gene evolution (Vakirlis et al., 2020). Our results therefore indicate that similar mechanisms may be at play in in *Triticeae*, and that they might have evolved lineage-specific regulatory networks, that induce the secretion of proteins in response to cold stress and/or different specialized fungal pathogens.

Interestingly, the majority of the identified apoplastic TSGs, have similarity to previously described *Triticeae*-specific *Lt-Cor* (low-temperature or cold-responsive) genes, that are involved in cold tolerance in wheat (Ohno *et al*., 2001). Some are known to be bifunctional as they also show antifungal activity against cold-adapted plant pathogens such as snow molds and stripe rust (Kuwabara and Imai, 2009; Gaudet *et al*., 2011). The *Triticum aestivum cold-regulated 7 (Tacr7)* gene family, for example, was shown to be up-regulated in wheat lines containing the leaf rust resistance genes *Lr34* (Hulbert *et al*., 2007). Indeed, we identified 13 apoplastic TSGs that have homology to the *Tacr7* gene family and that show significant up-regulation in both cold stressed and stripe rust infected wheat plants.

Additionally, we identified novel transmembrane proteins that have similarity with the *Wheat-Induced Resistance 1* (*WIR1*) gene family. As demonstrated by our transcriptomic analysis, members of this gene family are induced by a broad range of plant pathogens. In particular, *TaWIR1* (also found in our survey) inhibits the secondary hyphae growth of the non-adapted pathogen *Magnaporthe oryzae* (Tufan *et al*., 2012), and it was proposed to play an important role in non-host resistance. Finally, we showed that, compared to apoplastic TSGs, membrane-localized TSGs are under stronger diversifying selection. This is reminiscent of the co-evolutionary arms race between plant and pathogens, and further indicates the role of novel transmembrane proteins in the regulation of biotic stress responses.

To summarize, we identified *Triticeae*-specific genes that were previously described to be involved in *Triticeae* responses to cold and fungal pathogens. The fact that we were able to identify multiple previously undescribed *WIR1* and *Lt-Cor*-like genes, validates our approach of TSG identification and suggests that several candidate TSGs might be involved in important *Triticeae* adaptations. Importantly, since homology is restricted to the signal peptide and structural amino acids such as Cysteine and Proline, sequence similarity was too weak to be found by simple homology search. Only once we classified the candidate TSGs into families and we quantified their transcriptional response to stresses, similarities with *Lt-Cor* and *WIR* proteins became apparent. We propose that these TSGs identified here are interesting candidates for future functional studies on non-host resistance and response to environmental stress in *Triticeae*.

Three main mechanisms have been described for the evolution of lineage-specific genes from TEs, namely exonization, domestication of TE proteins, and capture and rearrangement of gene fragments (Jin *et al*., 2019). However, it is still unclear to what extent do TEs contribute to the evolution of *de novo* genes, and what molecular mechanisms underlie this process. In total, ∼18% (1,079 out of 5,942) of our candidate TSGs have homology to TEs at the DNA level. Only 12 of them showed significant protein sequence homology and 67 are found in alternative frames of TE proteins, suggesting that *de novo* evolution from non-coding sequences, rather than mere domestication of existing TE proteins, might be the main path for the evolution of TE-derived genes in *Triticeae*. Similar results were found in rice where most TE-derived lineage-specific genes were shown to originate *de novo* through rapid sequence divergence (Jin *et al*., 2019). In contrast, 93% of TE-derived lineage-specific genes in primates were shown to be exonized from *SINE* retrotransposons, which contain potential splice sites (Krull et al. 2005).

In total 764 (∼71%) of TE-derived TSGs described here were derived from LTR-retrotransposons of the *Gypsy* superfamily (Suppl. table S4). The next most represented TE superfamilies are *CACTA* transposons (122 genes) and *Copia* retrotransposons (83 genes). These results are consistent with previous findings (Sun et al. 2015; Jin et al. 2019) and highlight the importance of LTR retrotransposons and *CACTA* elements in the evolution of *de novo* genes in *Triticeae*.

Due to the rapid evolution of intergenic and non-coding sequences in *Triticeae*, it is very difficult to trace the evolution of a particular *de novo* gene to its very origins. However, we were able to largely resolve the origin of the *CACTA*-derived gene family TE2. Since all copies of the *CACTA* element (*DTC_clifford* family) from wheat and barley contained the complete CDS and only fragments of the CDS were found at different loci in the oat genome (none of them was associated to TEs), we propose that *CACTA* transposon in the *Triticeae* ancestor captured two non-coding, single-copy sequences into its promoter region. This proto-gene had the good “fortune” to be inside a highly proliferating *CACTA* element, as it could multiply, diversify and be transcribed along with the TE, until a new protein-coding transcript could emerge. In support of this hypothesis, previous studies have shown that DNA transposons can create lineage-specific genes through recombination of captured sequences (Jiang *et al*., 2004; Lai *et al*., 2005) and *CACTA*s have been correlated to recent gene duplications in wheat (Daron *et al*., 2014). Gene duplication has long been known to be important for adaptive evolutionary novelties (Panchy et al. 2016). Finally, all members of this gene family are derived from an AT-rich non-coding region and encode a N-terminal TM domain, further suggesting that the presence of a signaling domain, might be important for the selection of the newly evolved gene.

Approximately 93% of TE-derived *de novo* genes are localized inside or near regulatory regions such as long terminal repeats (LTRs) or terminal inverted repeats (TIRs). We found this subset of TSGs coming from TE non-coding regions being of particular interest. In fact, their GC content was particularly low, they were enriched in TM-encoding domains, and despite the majority of them (96%) being low-confidence genes, their expression profile in wheat (total number of expressed genes and average expression value) was similar to that of high-confidence genes. Vakirlis *et al*. (2020) showed that intergenic thymine-rich regions are hotspots for the emergence of adaptive membrane proteins, and since TE regulatory regions are usually AT-rich, such derived *de novo* proteins are therefore likely to contain transmembrane domains. Taken together, our results suggest that highly conserved AT-rich motifs might be functionally innovated into TM-domain encoding proteins and that TEs might be an important source of novel adaptive transmembrane proteins in *Triticeae*.

To conclude, in our study we developed an approach for the identification and characterization of *de novo* genes in *Triticeae*. We identified a total of 5,942 candidate *Triticeae*-specific of genes (TSGs), of which ∼60% were validated as protein-coding genes in wheat. This not only supports our approach but suggest that *de novo* gene formation is relatively rare and that *Triticeae* may possess approximately 779 lineage-specific genes per haploid genome. We demonstrated that 50% of stress-upregulated TSGs encode signaling sequences, the majority being related to important gene families involved in *Triticeae* adaptations to cold and fungal pathogens. This highlights the importance/need of novel secreted proteins in *Triticeae* stress adaptations. Finally, we show that TEs are responsible for ∼18% of our candidate TSGs, and we propose that functional innovation of TM domains from TE non-coding regions might be an important mechanism by which novel ORFs are selected and consequently evolve into adaptive stress-responsive TSGs.

## Acknowledgements

This work was supported by the Swiss National Foundation grant 31003A_163325.

## Author Contributions

MP and TW wrote and edited the manuscript, designed the study, and coordinated the research. MP and CP designed the pipeline. MP, TW, CP and AS performed bioinformatic analyses.

## Data Availability

The wheat genome annotation (IWGSC RefSeq v1.0) is openly available in the IWGSC Data Repository at https://wheat-urgi.versailles.inra.fr/Seq-Repository/Annotations. The rye genome annotation (Lo7 v1) is openly available via e!DAL at 10.5447/ipk/2020/29. The Barley genome annotation (IBSC v2) is openly available on the Ensembl Plants server (https://plants.ensembl.org/Hordeum_vulgare). The wheat gene expression values can be found on the Wheat Expression Browser at http://www.wheat-expression.com/download. Transcriptomic raw data are openly available on NCBI under bioproject codes PRJEB8798, PRJNA289545, PRJNA263755, PRJNA243835, PRJEB12497, PRJEB23056, PRJNA253535, PRJNA296894, PRJNA427159, and PRJNA327013.

