## Supplementary materials for "A survey of lineage-specific genes in *Triticeae* reveals *de novo* gene evolution from genomic raw material"

**Supporting information**

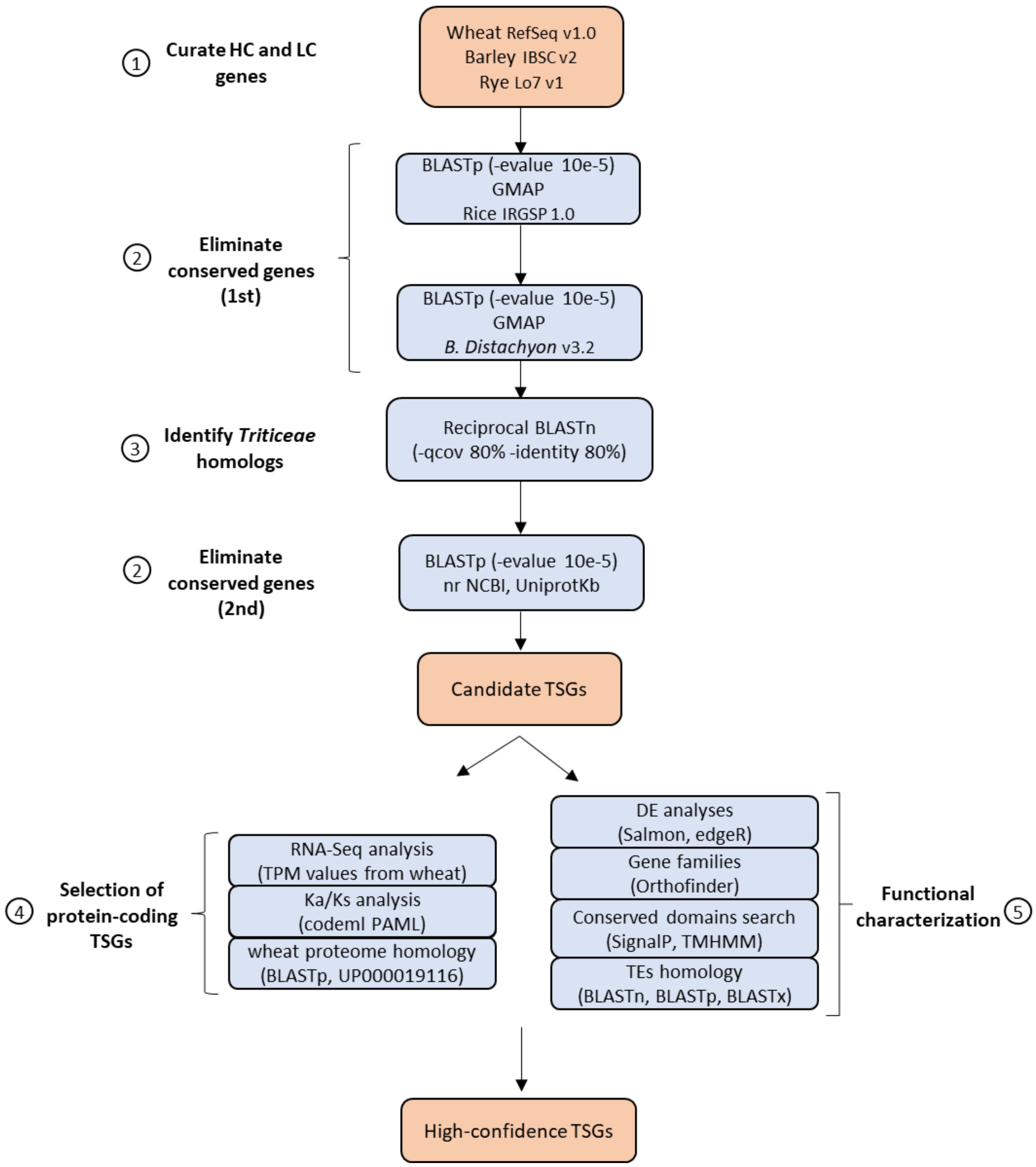


**Supplementary fig. S1. Pipeline for prediction of *Triticeae*-specific genes (TSGs).** As a first step, previously annotated high-confidence (HC) and low-confidence (LC) genes of *Triticeae* species were curated by filtering out all gene models with incomplete open reading frame (i.e., presence of in-frame stop codons and absence of start/stop codons). Then we selected candidate TSGs by searching for *Triticeae* genes that are annotated in at least two of the main *Triticeae* lineages (wheat, rye and barley) but have no homologs outside of the *Triticeae* clade (sequence homology searches against closely related outgroup species *B. distachion* and rice, and also against nrNCBI and UniprotKb databases). Then, we removed potential prediction artifacts by measuring the protein-coding potential of candidate TSGs. Genes with experimental evidence of translation and/or evidence of both transcription and negative selection were considered as “real” protein-coding TSGs. In addition, candidate TSGs were further functionally characterized by clustering gene families, analysing differential gene expression in wheat, predicting conserved domains (signal peptides and transmembrane domains), and investigating possible mechanisms of evolution from transposable elements.

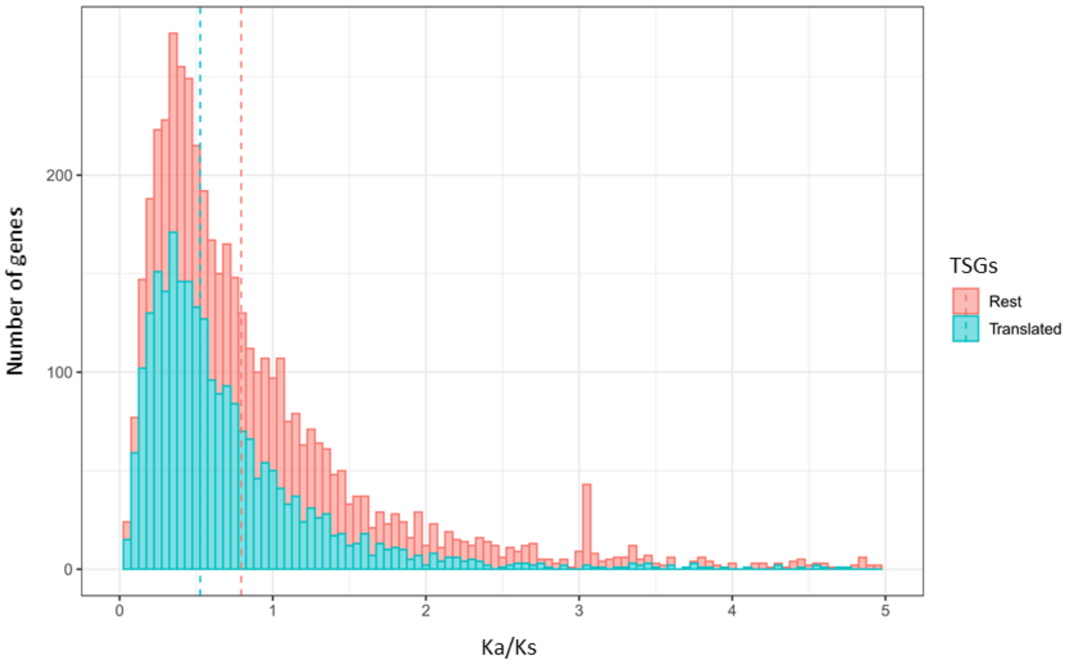


**Supplementary fig. S2. Barplot showing the distribution of Ka/Ks values of candidate *Triticeae* specific genes (TSGs).** Homologous TSG sequences between wheat, rye, and barley were compared at the protein level and the Ka/Ks values were calculated using the codeml program of the PAML package (Yang, 2007). Dashed lines indicate the median Ka/Ks values of 1,890 wheat TSGs that are supported by proteomic evidence (median Ka/Ks = 0.53; light blue) and of the remaining 4,052 TSGs (median Ka/Ks = 0.79; red). The median Ka/Ks value of translated wheat TSGs was chosen as cut-off for selecting TSGs that are under negative selection pressure. In total, we identified 2,365 TSGs that fall below this cut-off.

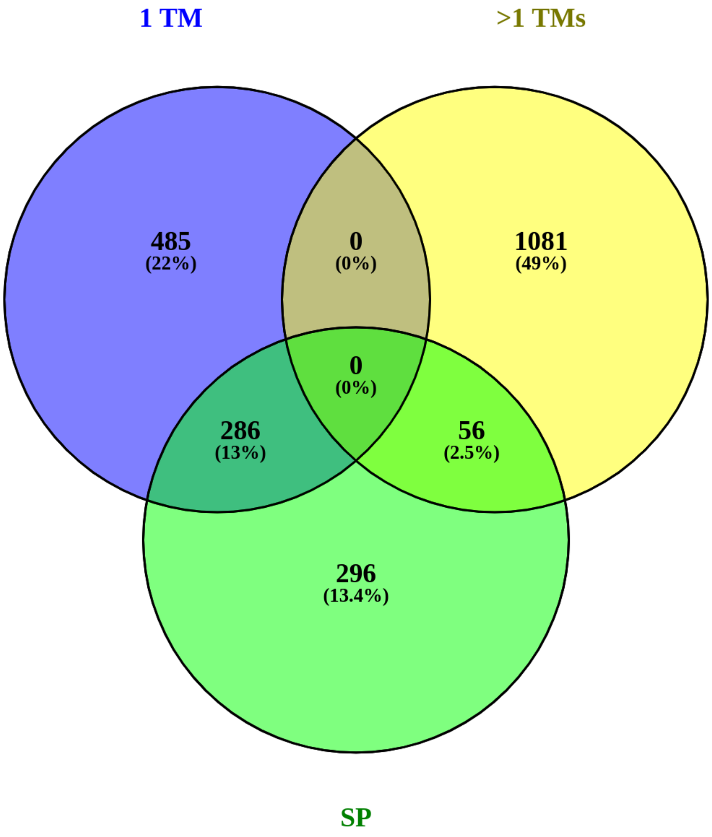


**Supplementary fig. S3. Manual curation of *Triticeae* specific genes (TSGs) encoding signaling sequences**. Because of the similarities between transmembrane domains (TMs) and signal peptides (SPs), the prediction of the cellular localization of proteins belonging to the secretory pathways can be ambiguous. To determine, which proteins are likely to be anchored to the plasma membrane and which ones might be secreted into the apoplast, we arbitrarily defined the following thresholds. Transmembrane proteins are characterized by i) at least one TM domain and no SP, ii) one SP and multiple TM domains or iii) low SP prediction value (<0.5). Apoplastic proteins are characterized by i) one SP or ii) one SP and one TM domain. SignalP (v5.0) and TMHMM (v2.0) were used to annotate SPs and TM domains, respectively.

**Supplementary table S1. Assessment of the integrity of high-confidence (HC) and low-confidence (LC) genes of *Triticeae* species (wheat, barley and rye).** Genes with incomplete (absence of terminal start and stop codons) and truncated (presence of in-frame stop codons) ORFs, might represent prediction artifacts. In order to better compare the gene content of rye and barley (diploids) to wheat (hexaploid), we indicate the number of genes for each wheat subgenome (A, B, D), including the "unknown" (Un) chromosome. Approximately 34,000-43,000 HC genes were annotated per haploid genome. Note that barley have a much larger number of incomplete ORFs, probably reflecting the improved gene annotation pipeline that was used for wheat and rye (IWGSC *et al.*, 2018).

| | | | | **Wheat subgenomes** | | | |
|---|---|---|---|---|---|---|---|
| **Gene Category** | **Rye** | **Barley** | **Wheat** | **A** | **B** | **D** | **Un** |
| HC genes | 34,441 | 42,732 | 110,790 | 36,302 | 36,738 | 35,021 | 2,728 |
| HC intact ORFs | 34,441 | 19,584 | 108,495 | 35,539 | 35,984 | 34,304 | 2,668 |
| HC no start | 0 | 19,162 | 0 | 0 | 0 | 0 | 0 |
| HC no stop | 0 | 7,891 | 2,294 | 763 | 754 | 717 | 60 |
| LC genes | 22,781 | 45,162 | 158,793 | 50,677 | 57,319 | 44,103 | 6,692 |
| LC intact ORFs | 20,400 | 18,964 | 111,478 | 35,741 | 40,718 | 30,967 | 4,051 |
| LC no start | 1,244 | 20,009 | 23,022 | 7,236 | 7,983 | 6,448 | 1,355 |
| LC no stop | 1,642 | 12,991 | 19,883 | 6,242 | 6,804 | 5,443 | 1,394 |
| LC in-frame stop | 0 | 0 | 17,416 | 5,682 | 6,406 | 4,666 | 662 |

**Supplementary table S2. Summary of RNA-Seq datasets used for differential expression (DE) analyses.** Note that, since different wheat lines, plant tissues and developmental stages were considered, the total number of DE genes identified in different experiments should be compared with caution.

| Wheat line(s) | Treatment | Tissue | Dev. Stage | Time points | SRA accession |
|---|---|---|---|---|---|
| Drifter | *Z. tritici* | leaf | NA | 3,7,12,14,28 dpi | SRP077418 |
| Chinese Spring | *B. graminis* | leaf | seedling | 2, 7 dpi | PRJEB23548, PRJNA553193 |
| NA | *B. graminis* | leaf | seedling | 24,48,72 hpi | SRP041017 |
| NA | *P. Striiformis* | leaf | seedling | 24,48,72 hpi | SRP041017 |
| Vuka, Avocet+Yr5 | *P. Striiformis* | leaf | 3 leaf stage | 1,2,3,5,7,9,11 dpi | ERP013983 |
| NIL51 from BC55F2 Remus in CM-82036 | *F. graminareum* | spike | anthesis | 3,6,12,24,48 hpi | ERP013983 |
| Janz*2 NIL1 susceptible, Janz*2 NIL1 resistant | *F. pseudograminearum* | shoot | NA | 5 dpi | SRP048912 |
| Chinese Spring | Chitin | leaf | 3 leaf stage | NA | PRJEB23056 |
| Chinese Spring | Flagellin 22 | leaf | 3 leaf stage | NA | PRJEB23056 |
| Manitou | Cold (4°C) | shoot | 3 leaf stage | 2 weeks | SRP043554 |

**Supplementary table S3. Transcriptomic analysis of two *Triticeae*-specific gene (TSG) families showing homology to wheat mitochondrial DNA.** Salmon was used for mapping and quantifying the number of mapped RNA-Seq reads against the wheat CDS. The number of mapped reads (NumReads) were used for differential expression analysis using the R package edgeR (see methods). Average RPKM values were calculated from three biological replicates. Samples were compared to the control (3dpi) in a pairwise manner, and the likelihood ratio test was used for testing for differential gene expression. Only genes with a log2FC > |1.5| and an adjusted p-value (FDR) < 0.01 were considered as differentially expressed. LogFC values represent the change in gene expression at 28 days post infection (dpi) in comparison to 3 dpi. Note that even though these TSGs show homology to the mitochondrial DNA, none of them have mitochondrial targeting peptides, thus suggesting that, after the transfer to the nuclear genome, these putative novel genes arose from DNA of mitochondrial origin but not directly from mitochondrial CDS. Significant up-regulation in response to *Z. tritici* infection (especially at 28dpi) suggests that these genes might have acquired a novel function in the wheat immune response to *Z. tritici*.

| | | | Average RPKM | | | | | | |
|---|---|---|---|---|---|---|---|---|---|
| **Wheat TSG** | **TSG homolog** | **Ka/Ks** | **3dpi** | **7dpi** | **12dpi** | **14dpi** | **28dpi** | **logFC_28dpi** | **FDR** |
| TraesCS2B01G920500LC | HORVU0Hr1G019860L | 0.000 | 0.038 | 0.000 | 0.000 | 4.522 | 15.735 | NA | NA |
| TraesCS5B01G152200LC | HORVU0Hr1G019860L | 0.000 | 0.038 | 0.000 | 0.000 | 0.874 | 160.432 | 11.353 | 4.71E-04 |
| TraesCS5B01G153700LC | HORVU0Hr1G019860L | 0.449 | 0.038 | 0.000 | 0.000 | 0.616 | 0.000 | NA | NA |
| TraesCS5B01G531200LC | HORVU0Hr1G019860L | 0.000 | 0.038 | 0.000 | 0.000 | 0.616 | 160.432 | 11.339 | 2.08E-03 |
| TraesCSU01G250300LC | HORVU0Hr1G019860L | 0.000 | 0.038 | 0.000 | 0.000 | 0.616 | 160.432 | 11.339 | 2.08E-03 |
| TraesCS2B01G406900LC | SECCE2Rv1G0087220 | 0.758 | 0.000 | 0.000 | 0.000 | 1.407 | 212.673 | 12.758 | NA |
| TraesCS2D01G255900LC | SECCE2Rv1G0087220 | 0.000 | 0.000 | 0.000 | 0.000 | 0.000 | 0.000 | NA | 4.08E-06 |
| TraesCS5A01G200700LC | SECCE2Rv1G0087220 | 0.000 | 0.114 | 0.035 | 0.078 | 3.648 | 387.460 | 11.340 | 1.78E-16 |
| TraesCS5B01G152300LC | SECCE2Rv1G0087220 | 0.000 | 0.136 | 0.045 | 0.000 | 3.607 | 322.093 | 10.748 | 1.31E-06 |
| TraesCS7B01G819400LC | SECCE2Rv1G0087220 | 0.440 | 0.024 | 0.000 | 0.000 | 0.080 | 0.000 | NA | NA |
| TraesCSU01G624500LC | SECCE2Rv1G0087220 | 0.000 | 0.136 | 0.045 | 0.000 | 3.607 | 322.093 | 10.748 | 1.31E-06 |

**Supplementary table S4. Triticeae-specific genes (TSGs) related to transposable elements (TEs).** The table summarize the number of TSGs (for each *Triticeae* species) that are associated to specific TE superfamilies. Note that retrotransposons of the *Gypsy* superfamily (RLG) contribute to the vast majority (764 out of 1,079; ~71%) of TE-derived TSGs, followed by *CACTA* DNA transposons (DTC; ~11%) and *COPIA* retrotransposons (RLC; ~8%). Interestingly, we see that TSGs belonging to *Gypsy* and *CACTA* superfamilies are especially abundant in wheat.

| TE Superfamily | Wheat | Barley | Rye | Total |
|---|---|---|---|---|
| RLG | 662 | 73 | 29 | 764 |
| DTC | 96 | 19 | 7 | 122 |
| RLC | 35 | 21 | 27 | 83 |
| RLX | 50 | 0 | 8 | 58 |
| DTH | 26 | 1 | 1 | 28 |
| RSX | 5 | 4 | 1 | 10 |
| DTM | 1 | 1 | 4 | 6 |
| DHH | 1 | 1 | 1 | 3 |
| RIX | 3 | 0 | 0 | 3 |
| DTT | 1 | 1 | 0 | 2 |

**Supplementary table S5. Analysis of most abundant *Triticeae*-specific gene families.** Families containing ≥ 10 gene members were further analysed for presence of signaling motifs, such as transmembrane domains (TM) and signal peptides (SP), and also for sequence homology to transposable elements (TEs). Interestingly, 14 out of 35 gene families, including the 3 largest ones, contain genes encoding for TM domains. This suggests that the presence of TM domains might have played an important role in the evolution and diversification of *de novo* genes. In addition, 5 out of 7 gene families deriving from TEs also encode TM domains, suggesting a correlation between TEs and TM-encoding regions.

| | Number of family members | | | | Conserved domains | | |
|---|---|---|---|---|---|---|---|
| **Family** | **Wheat** | **Barley** | **Rye** | **Total** | **TM** | **SP** | **TE homology** |
| OG0000000 | 12 | 2 | 134 | 148 | TM | NA | NA |
| OG0000001 | 13 | 2 | 128 | 143 | TM | NA | NA |
| OG0000002 | 73 | 2 | 1 | 76 | TM | SP | RLG_Scer_Sabrina_D1_consensus-3 |
| OG0000004 | 25 | 2 | 20 | 47 | NA | NA | NA |
| OG0000005 | 44 | 1 | NA | 45 | NA | NA | RLG_Taes_Egug_consensus-1 |
| OG0000007 | 26 | 1 | 15 | 42 | NA | NA | NA |
| OG0000009 | 1 | NA | 33 | 34 | NA | NA | NA |
| OG0000010 | 2 | 1 | 28 | 31 | TM | NA | NA |
| OG0000011 | 30 | NA | 1 | 31 | TM | NA | NA |
| OG0000012 | 21 | 1 | 7 | 29 | NA | NA | NA |
| OG0000013 | 24 | 3 | 1 | 28 | TM | NA | NA |
| OG0000014 | 21 | 2 | 4 | 27 | TM | NA | NA |
| OG0000015 | 20 | 1 | 4 | 25 | NA | NA | NA |
| OG0000018 | 21 | NA | 1 | 22 | TM | NA | DTC_Atau_Jude_AF446141-1 |
| OG0000019 | 17 | 1 | 1 | 19 | TM | NA | RLG_Hvul_Sabrina_B_consensus-1 |
| OG0000017 | 15 | 1 | 1 | 17 | NA | NA | NA |
| OG0000020 | 16 | 1 | NA | 17 | TM | NA | NA |
| OG0000021 | 15 | 1 | 1 | 17 | TM | NA | RLG_Scer_Sabrina_D1_consensus-3 |
| OG0000022 | 11 | NA | 6 | 17 | NA | NA | NA |
| OG0000023 | 1 | NA | 15 | 16 | NA | NA | NA |
| OG0000024 | 14 | NA | 2 | 16 | NA | NA | NA |
| OG0000026 | 3 | NA | 12 | 15 | NA | NA | NA |
| OG0000027 | 12 | NA | 3 | 15 | NA | NA | NA |
| OG0000029 | 9 | 1 | 4 | 14 | NA | NA | NA |
| OG0000028 | 9 | 2 | 1 | 12 | NA | NA | NA |
| OG0000030 | 9 | 1 | 2 | 12 | NA | NA | NA |
| OG0000031 | 9 | 2 | 1 | 12 | NA | NA | RLG_Hvul_BAGY2_consensus-2 |
| OG0000034 | 5 | 4 | 2 | 11 | NA | NA | NA |
| OG0000036 | 7 | 1 | 3 | 11 | TM | SP | NA |
| OG0000038 | 2 | 1 | 7 | 10 | TM | SP | NA |
| OG0000039 | 9 | 1 | NA | 10 | TM | NA | RLG_Scer_Sabrina_A2_consensus-3 |
| OG0000040 | 8 | 1 | 1 | 10 | NA | SP | NA |
| OG0000041 | 5 | 1 | 4 | 10 | NA | NA | NA |
| OG0000042 | 1 | NA | 9 | 10 | NA | NA | NA |
| OG0000044 | 1 | NA | 9 | 10 | NA | NA | NA |
